# Chromosome dynamics in bacteria: triggering replication at opposite location and segregation in opposite direction

**DOI:** 10.1101/521781

**Authors:** Ady B. Meléndez, Inoka P. Menikpurage, Paola E. Mera

**Affiliations:** Department of Chemistry and Biochemistry, New Mexico State University, Las Cruces, NM 88003, USA

**Keywords:** Chromosome replication, chromosome segregation, DnaA, ParA, centromere, *Caulobacter crescentus*

## Abstract

The accurate onset of chromosome replication and segregation are fundamental for the survival of the cell. In bacteria, regulation of chromosome replication lies primarily at the initiation step. The bacterial replication initiator DnaA recognizes the origin of replication (*ori*) and opens this double stranded site allowing for the assembly of the DNA replication machinery. Following the onset of replication initiation, the partitioning protein ParA triggers the onset of chromosome segregation by direct interactions with ParB-bound to the centromere. The subcellular organization of *ori* and centromere are maintained after the completion of each cell cycle. It remains unclear what triggers the onset of these key chromosome regulators DnaA and ParA. One potential scenario is that the microenvironment of where the onset of replication and segregation take place hosts the regulators that trigger the activity of DnaA and ParA. In order to address this, we analyzed whether the activity of DnaA and ParA are restricted to only one site within the cell. In non-dividing cells of the alpha proteobacterium *Caulobacter crescentus*, *ori* and centromere are found near the stalked pole. To test DnaA’s ability to initiate replication away from the stalked pole, we engineered a strain where movement of *ori* was induced in the absence of chromosome replication. Our data show that DnaA can initiate replication of the chromosome independently of the subcellular localization of *ori*. Furthermore, we discovered that the partitioning protein ParA was functional and could segregate the replicated centromere in the opposite direction from the new pole toward the stalked pole. We showed that the organization of the ParA gradient can be completely reconstructed in the opposite orientation by rearranging the location of the centromere. Our data reveal the high flexibility of the machineries that trigger the onset of chromosome replication and segregation in bacteria. Our work also provides insights into the coordination between replication and segregation with the cellular organization of specific chromosomal loci.

## INTRODUCTION

Triggering the onset of chromosome replication at the correct frequency and under the right conditions is central to the survival and proliferation of the cell. In bacteria, DnaA is the highly conserved initiator of chromosome replication [1]. DnaA is an AAA+ protein that recognizes a specific chromosomal locus known as the origin of replication (*ori*) [2, 3]. DnaA binds *ori* forming a helical right-handed polymeric structure, which exerts a torsional stress that results in the opening of the DNA strands [4, 5]. Once DnaA opens the double stranded chromosome, the replication machinery assembles at *ori* and initiates chromosome replication bidirectionally.

The multiple levels of DnaA’s regulation reveal the significant investment the cell makes to accurately initiate chromosome replication. DnaA’s ability to bind *ori* can be regulated by either *ori*-binding proteins that outcompete DnaA for binding sites, or by non-*ori* chromosomal loci that can sequester DnaA away from *ori* [6, 7]. Proteins like DiaA (*E. coli*) and HobA (*H. pylori*) facilitate DnaA’s ability to form a helical polymeric structure at *ori* by recruiting multiple DnaA molecules [8–10]. Furthermore, the ability of DnaA to open the DNA strands can be turned off by the induction of its ATPase activity. Only DnaA bound to ATP can form the polymeric structures at *ori* required for strand opening [11]. DnaA’s activity can be turned off by HdaA, which stimulates DnaA-ATP hydrolysis [12–14]. In vitro analyses of DnaA’s interaction with membrane phospholipids have also been shown to induce the dissociation of the nucleotide bound and thus potentially regulate DnaA’s activity [15, 16]. In *C. crescentus,* the levels of DnaA in the cell can also be cell-cycle regulated [17–20].

The subcellular location where DnaA initiates chromosome replication is established by the position of *ori* inside the cell. Depending on the bacterial species, the subcellular location of *ori* varies significantly. For instance, *C. crescentus ori* is found at one pole (the stalked pole), whereas in *E. coli* and *B. subtilis*, *ori* is found near mid-cell. However, within a single species, the subcellular location of *ori* is strictly retained at the same position in non-dividing cells and re-established soon after chromosome replication and segregation initiate in actively dividing cells. In non-dividing cells of *C. crescentus*, *ori* is retained near the stalked pole by the interaction between the anchoring protein PopZ and the centromere (*parS)* [21–23]. Thus, DnaA initiates chromosome replication at ori near the stalked pole and terminates replication at *ter* locus near the opposite pole (referred here as the new pole) (Fig. 1).

**Figure 1.**
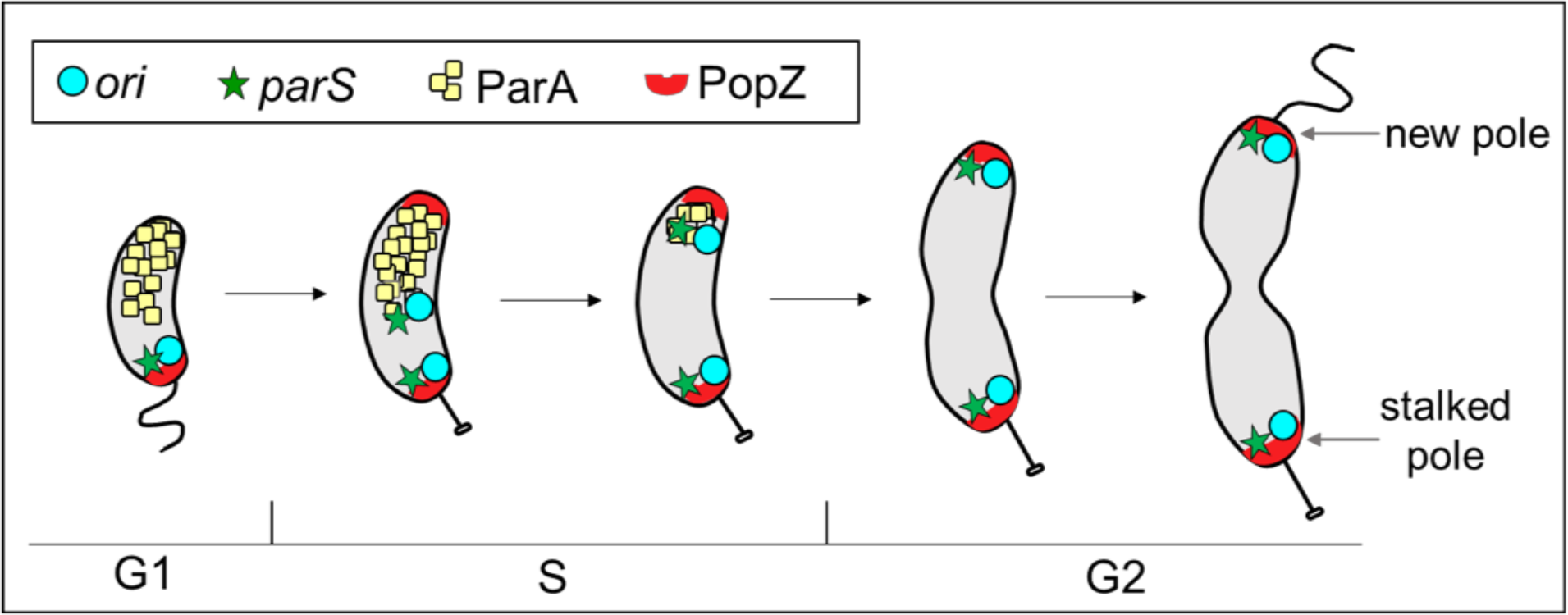
Cell-cycle dependent dynamics of *Caulobacter crescentus.* Localization of two chromosomal foci [origin of replication (*ori)* and centromere (*parS*)] and two proteins involved in chromosome segregation [ParA and PopZ] over the course of a normal cell cycle. Non-dividing cells have *ori* (cyan) and *parS* (green) localized near the stalked pole. PopZ anchors the centromere region *parS* and the *parS*-binding protein ParB complex at the stalked pole. Once replication initiates, two foci of *ori* and two foci of *parS* are observed. Upon the onset of chromosome replication and segregation, a second PopZ foci appears at the new pole. The new duplicated *ori* and *parS* are the first regions to move in a ParA-dependent manner to the new pole.

Once chromosome replication initiates at *ori*, the replication fork must pass through the centromere (*parS*, 8 kilobases away from *ori*) for chromosome segregation to initiate [24]. The centromere is bound by a multi-protein complex that plays essential roles in the cell cycle [21–25]. For example, the chromosome is anchored at one pole by interactions between the protein ParB and PopZ (a polar anchoring protein) [21–23] (Fig. 1). Soon after replication initiates, one of the two ParB-coated centromeres is segregated to the new pole by direct interactions between ParB and the ATPase ParA [24, 26–29]. ParA forms a stable gradient with concentrations gradually decreasing from the new pole to the stalked pole. The establishment of this ParA gradient is required for transport directionality [30]. ParB induces ATP-hydrolysis of ParA, releasing ParA from this gradient. Notably, ParA’s stable gradient is known to be established well before chromosome replication and segregation are initiated [26–28]. Thus, the mechanism that triggers ParA to initiate segregation of one centromere to the opposite pole remains unclear.

What regulates the timing for DnaA to initiate replication with such precise periodicity remains unsolved. What regulates the timing for ParA to initiate segregation only after the replication of the centromere is completed also remains unsolved. One potential scenario is that the microenvironment of the stalked pole in *C. crescentus*, where the onset of replication and segregation take place, hosts the regulators that trigger the activity of DnaA and ParA. With the goal to further understand what regulates the onset of these key cell cycle regulators (DnaA and ParA), we examined whether the onset of replication and segregation are restricted to *ori*-centromere’s intrinsic localization, which in *C. crescentus* is near the stalked pole. To resolve this question, we genetically engineered a *C. crescentus* strain where movement of *ori/*centromere can be triggered in the absence of replication initiation [31]. Once *ori* was translocated to different sites over the cell, we tested for competency of replication initiation and chromosome segregation. Our data revealed that DnaA and ParA are capable of initiating chromosome replication and segregation irrespective of the cellular location of *ori* and centromere. We show that the organization of the ParA gradient is completely flipped in orientation once the un-replicated centromere localized at the new pole. Our results uncover the robustness and flexibility of the chromosome replication and segregation machineries in bacteria.

## RESULTS

### Construction of indicator strain with fluorescently labeled origin of replication

To determine the cellular localization of *ori*, we engineered a fluorescent tag to be inserted near *ori* using the *Yersinia pestis parS*(pMT1) chromosomal sequence and its corresponding gene-encoding ParB(pMT1) fluorescently tagged [32]. The *parS*-ParB(pMT1) system from *Yersinia* has been previously used in *C. crescentus* and shown not to interfere with the activity of the native *C. crescentus* ParABS partitioning system [24]. Consistent with those findings, our analyses of growth curves and Colony Forming Units (CFU) revealed that strains with the *parS*(pMT1) insertion near *ori* and the expression of *parB*(pMT1) have no significant effect on the doubling time and viability compared to wild-type (Fig. S1. A & B). To control the expression of *dnaA*, we first engineered an additional copy of *dnaA* to replace the gene *vanA* resulting in the expression of *dnaA* regulated by the VanA promoter (*vanA::dnaA*) [33]. Using this merodiploid strain, the native gene encoding for DnaA was replaced with a spectinomycin cassette leaving no scars on the genome. The final indicator strain with a fluorescent label near *ori* and encoding a single copy of *dnaA* (regulated by P_*van*_) is referred here as PM500 with the genotype *xylX::parB*(pMT1), *parS*(pMT1) at nucleotide 1,108, *dnaA*::Ω, *vanA*::*dnaA*. For simplification from here on, we will refer to the *parS*(pMT1) localized near *ori* simply as *ori*.

### Sub-physiological levels of DnaA results in translocation of *ori* away from stalked pole

In *C. crescentus*, the centromere is the first chromosomal locus to segregate away from the stalked pole [24]. Previous analyses of cells expressing sub-physiological levels of DnaA (not sufficient to initiate replication) revealed a DnaA-dependent and replication-independent segregation of the centromere [31]. Under these sub-physiological levels of DnaA, cells move the un-replicated centromere from the stalked pole to the new pole. Based on those results, we asked if the same sub-physiological levels of DnaA could also trigger the movement of *ori*, independently of replication. To reach sub-physiological levels of DnaA in the cell, we used the same vanillate promoter to regulate *dnaA* expression [31]. Using our indicator strain PM500, we tracked the localization of *ori* using high-resolution microscopy. Our results show that sub-physiological levels of DnaA trigger the translocation of *ori* away from the stalked pole independently of chromosome replication (Fig. 2A). As the time of DnaA depletion increased, cells displayed un-replicated *ori* focus progressively moving toward the new pole (Fig. 2B). Within 3 h of DnaA depletion, ~ 75 % of cells exhibited their *ori* localized at the new pole. These results resemble the replication-independent movement of the centromere suggesting that *ori* and centromere translocate in a DnaA-dependent manner likely due to their close proximity on the chromosome [31].

**Figure 2.**
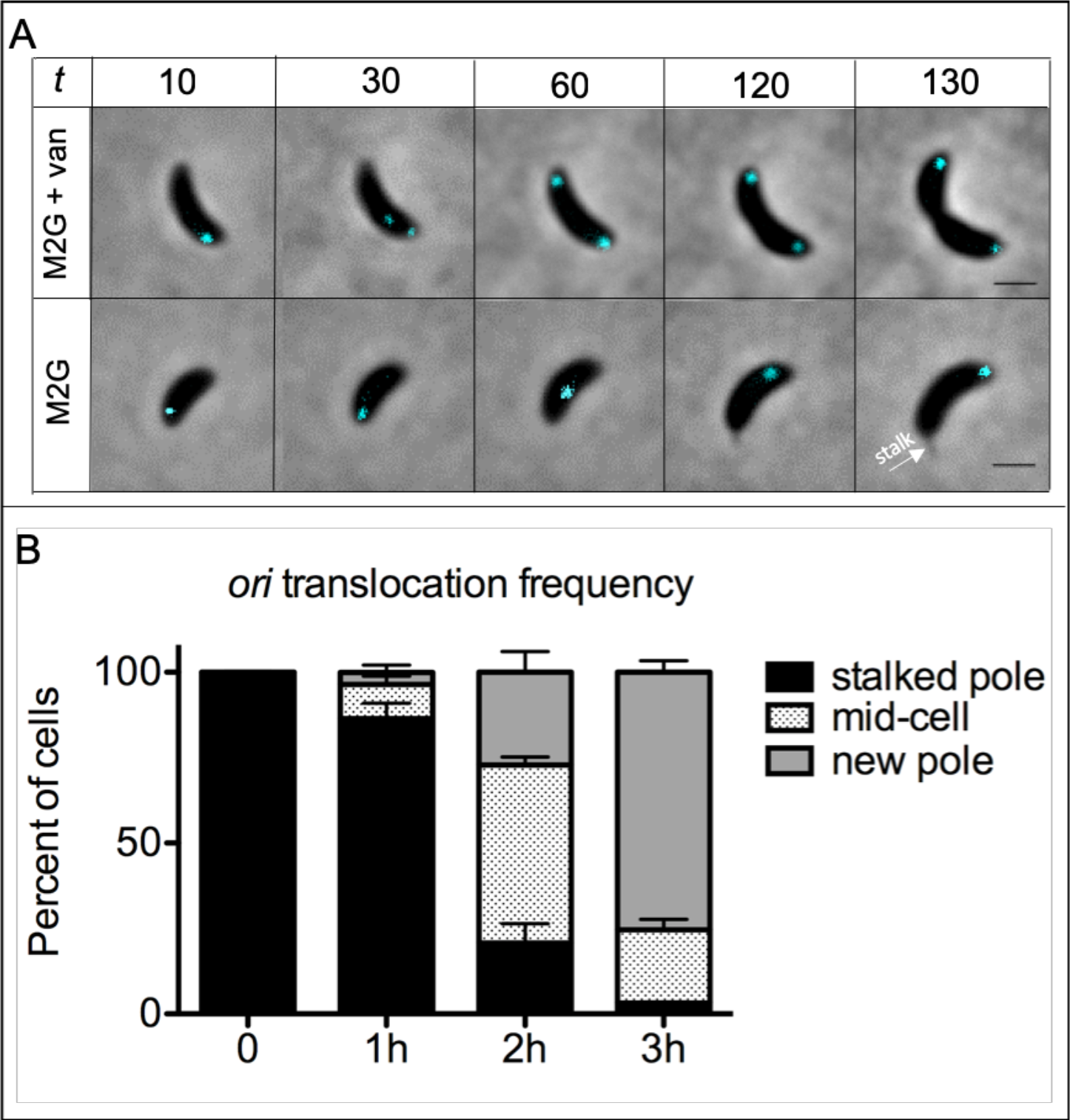
Translocation of *ori* triggered by DnaA in the absence of chromosome replication. (A) Time lapse of indicator strain [PM500: *parS*(pMT1), *vanA*::*dnaA*, Δ*dnaA*, pxylX::cfp- *parB*(pMT1)] with fluorescent tag near *ori* (~1kb) and DnaA expression regulated by the vanillate promoter P_VanA_. Cells grown in M2G with vanillate were synchronized and swarmer cells were spotted on 1 % agarose M2G pads in the presence (top row) or absence (bottom row) of vanillate (250uM). Cells were imaged with phase contrast and CFP-mediated fluorescence microscopy every 30 minutes. (B) Plotted are the mean ± SD percentages from three independent experiments of cells with translocated *ori* to the middle or new pole as time of DnaA depletion increases.

### DnaA’s ability to initiate replication is not restricted to the stalked pole

In *C. crescentus*, DnaA initiates replication only once per cell cycle and only in stalked cells, which have their *ori* localized at the stalked pole (Fig. 1). To determine whether DnaA can initiate replication outside the stalked pole, we analyzed cells that had undergone replication-independent translocation of *ori*. To track replication initiation, we supplemented vanillate to cells with *ori* localized outside the stalked pole and followed the appearance of two *ori* foci (Fig. 3A). Thirty minutes after the addition of vanillate, ~ 60 % of cells with *ori* localized at the new pole had initiated chromosome replication as evidenced by two clearly separated *ori* foci. Quantification of the frequencies of replication initiation based on location of *ori* after 3 h DnaA depletion revealed that replication initiation was triggered slightly quicker, yet statistically significant, from *ori* localized at the new pole compared to those localized at the stalked pole (Fig. 3B). Consistent with these results, a similar pattern in rates of replication initiation based on location of *ori* were observed when tracking the location of a chromosomal locus found 8 kilobases away from *ori* (Fig. S2). In sum, our results revealed that DnaA’s activity as a replication initiator is not restricted to the stalked pole.

**Figure 3.**
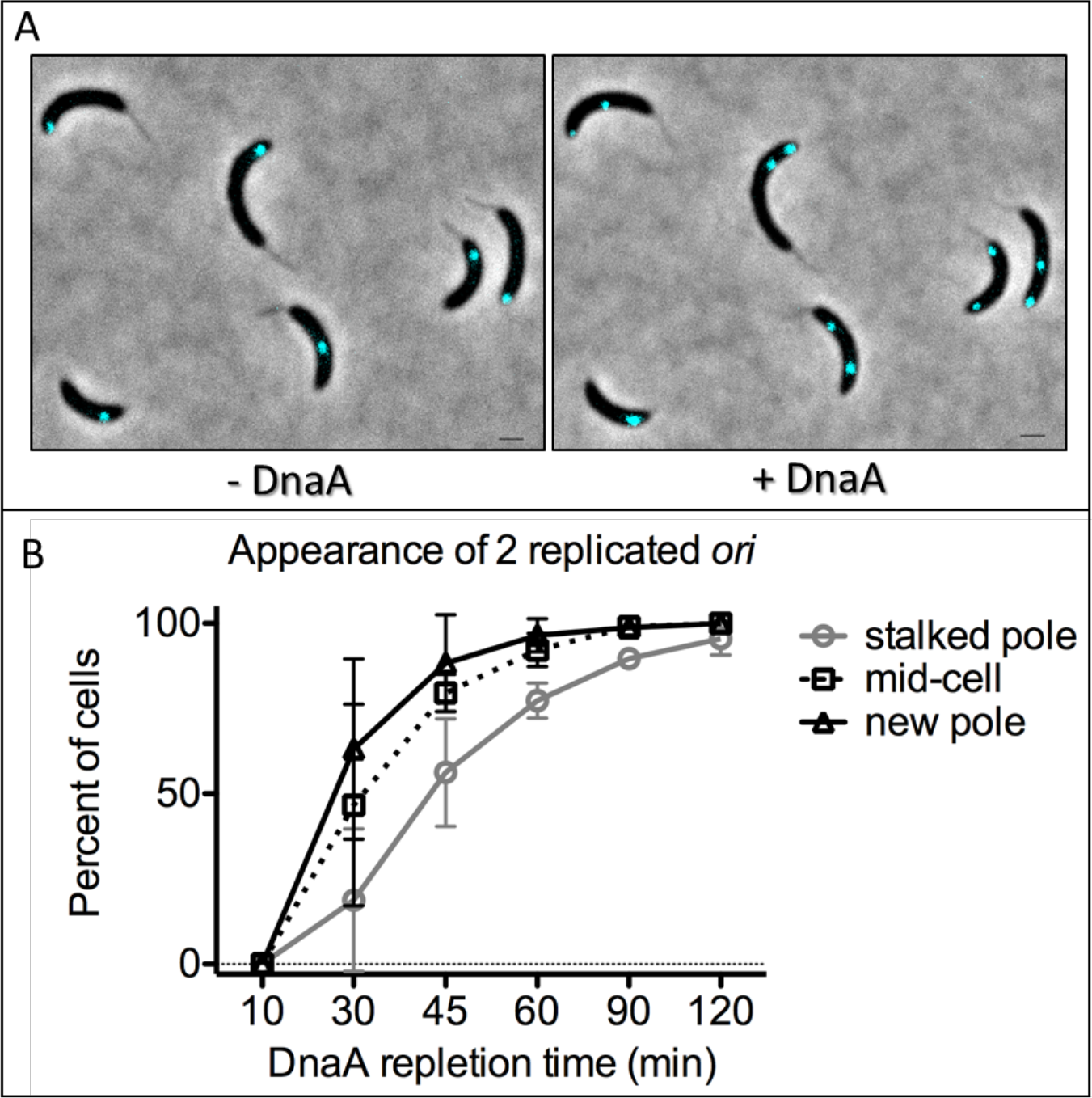
Chromosome replication is not limited to the stalked pole. A) Cells with fluorescent tag near *ori* [PM500, *parS*(pMT1) near *ori*, *vanA*::*dnaA*, Δ*dnaA*, *pxylX::cfp*-*parB* (pMT1)] were synchronized and DnaA was depleted by growing them in liquid M2G media without vanillate. At 3 h of DnaA depletion, un-replicated *ori* foci translocated to the opposite pole, middle or stayed at the stalked pole (left panel). After the depletion period, DnaA expression was induced by supplementation of vanillate (250 μM). Within 30 minutes of DnaA repletion, cells were able to initiate chromosome replication (right panel) as evidenced by two *ori* (cyan) foci. B) The appearance of two replicated *ori* foci were quantified from three independent fluorescence microscopy time-lapses. The plot represents the mean ± SD percent of cells that started replication at different time points. Analyses of two-way ANOVA in-between the frequencies of replication at stalk pole vs. new pole are significantly different at time points 30-min and 45-min; ***p<0.001 and *p<0.05.

### Centromeres are effectively segregated in the opposite direction

Based on our results that DnaA is able to initiate replication from the new pole (Fig. 3), we then formulated our next question concerning chromosome segregation. Can the partitioning system ParABS initiate segregation of the centromere from the new pole toward the stalked pole, which in this case would be in the opposite direction? To test this, we used a *C. crescentus* strain in which the native *parB* gene was replaced with the fusion gene encoding CFP-ParB and in which the only copy of *dnaA* was regulated under the vanillate promoter (PM109) [31]. In *C. crescentus*, the partitioning protein ParB binds directly to the centromere [24, 34]. Thus, we can track centromere movement by using cells expressing a functional fusion protein CFP-ParB. PM109 cells grown in the presence of vanillate were shown to display wild-type dynamics for CFP-ParB localization [31]. When the inducer vanillate is removed from the growth media of PM109, cells are exposed to sub-physiological levels of DnaA insufficient to initiate chromosome replication [31]. Fluorescent imaging of PM109 cells grown without vanillate revealed that the localization of a single CFP-ParB focus with the following distribution: ~ 37 % at or near the new pole, ~ 54 % at or near the stalked pole, and ~ 10 % at/around mid-cell (Fig. S3) These percent distribution of un-replicated centromere localization triggered in a replication-independent manner are consistent with previous analyzes [31].

To determine the ability of ParABS to trigger segregation of the centromere in the opposite direction, we tracked the movement of CFP-ParB in PM109 cells with translocated un-replicated centromeres supplemented with vanillate (Fig. 4). Similar to our analyses of *ori*, two clearly separated centromeres were observed soon after *dnaA* expression was reestablished, irrespective of the initial localization of centromere prior to *dnaA* induction. Notably, cells with two CFP-ParB foci were able to also segregate their centromeres to the cell poles, irrespective of the initial localization of centromere prior to *dnaA* induction (Fig. 4A). Quantification of these data revealed that the rate at which centromeres were segregated to the cell poles were significantly faster in cells with centromeres that departed from the mid-cell (Fig. 4 B & C). Chromosome segregation that initiated from either the stalked pole or the new pole was not statistically different. These data suggest that the partitioning protein ParA was able to reorganize its gradient in order to segregate centromeres from mid-cell or from the new pole.

**Figure 4.**
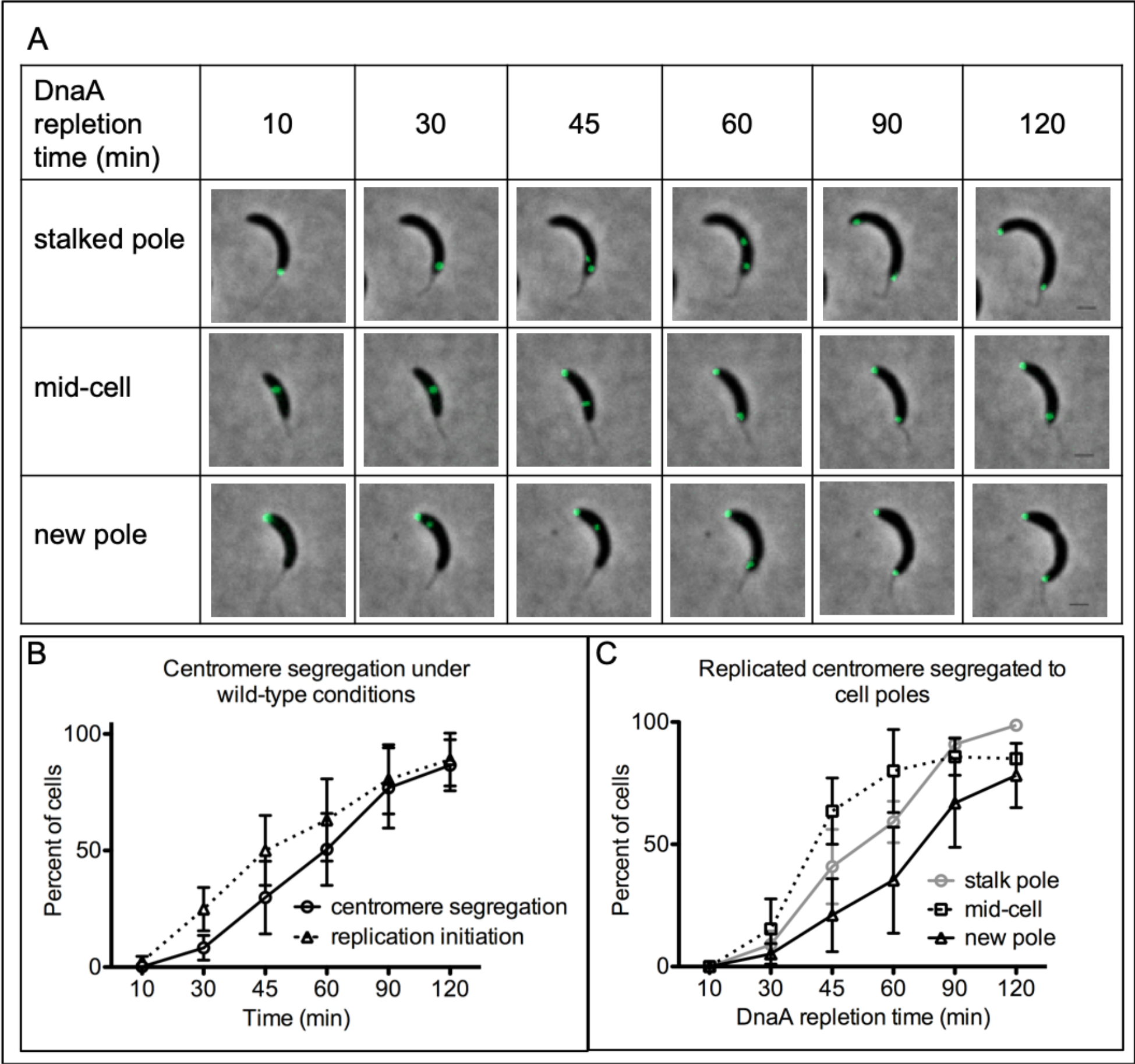
Translocated centromeres effectively segregate in opposite direction. (A) Time-lapse of centromere segregation (green foci represent CFP-ParB/*parS*) starting from the stalked pole, mid-cell or new pole of PM109 (*parB*::*cfp-parB*, *dnaA*::Ω, *vanA*::*dnaA*) cells after 3 h of DnaA depletion. Cells imaged were synchronized prior to DnaA depletion in M2G media (2 mL, OD_600_ ~ 0.1) and vanillate (250 μM) was added (time 0 min) to induce the expression of DnaA. 2 μL of cells were spotted on 1 % agarose pads supplemented with vanillate (250 μM). Scale bar = 1 μm. (B) Chromosome segregation of synchronized PM109 (*parB*::*cfp-parB*, *dnaA*::Ω, *vanA*::*dnaA*) cells under wild-type conditions (media supplemented with vanillate). (C) Plotted are the mean ± SD of cells with centromeres segregation to the cell poles based on initial localization of CFP-ParB/*parS*. Data represents three independent experiments. Statistical analyses of two-way ANOVA in-between the frequencies of segregation at mid-cell and new pole are significantly different at 45-min and 60-min time points, **p<0.01 and the frequencies of segregation at or near stalked pole and mid-cell at 45-min. *p<0.05.

### Active ParA is required for centromere segregation in the opposite direction

*C. crescentus* cells expressing ParA variants that are unable to hydrolyze ATP cannot segregate their centromere to the cell poles [24, 26, 35]. To determine whether the segregation of centromere segregation observed from the new pole/mid-cell is ParA-dependent, we tracked CFP-ParB localization in a merodiploid strain that encodes the wild-type allele of *parA* at the native locus and a dominant negative mutant *parA* allele regulated by the xylose-inducible promoter. This dominant negative allele encodes a missense mutation in the ATPase domain (ParA^D44A^) of ParA that inhibits chromosome segregation [36]. To test for segregation in the opposite direction, we first allowed cells to translocate their un-replicated centromeres by growing them in growth media devoid of vanillate. Xylose was then supplemented to the growth media 1 h prior to the addition of vanillate so that replication initiation was induced in the presence of the dominant negative ParA^D44A^. Our data revealed that > 80 % of cells expressing wild type ParA were able to segregate the centromeres to the poles, as evidenced by one CFP-ParB focus at each pole (Fig. 5). However, cells expressing the dominant negative ParA^D44A^ after replication initiated outside the stalked pole failed to segregate their replicated centromeres as evidenced by two CFP-ParB near each other (Fig. 5 A & B). These data strongly suggest that the segregation observed of the centromere in the opposite direction from the new pole to the stalked pole requires an active chromosome segregation machinery.

**Figure 5.**
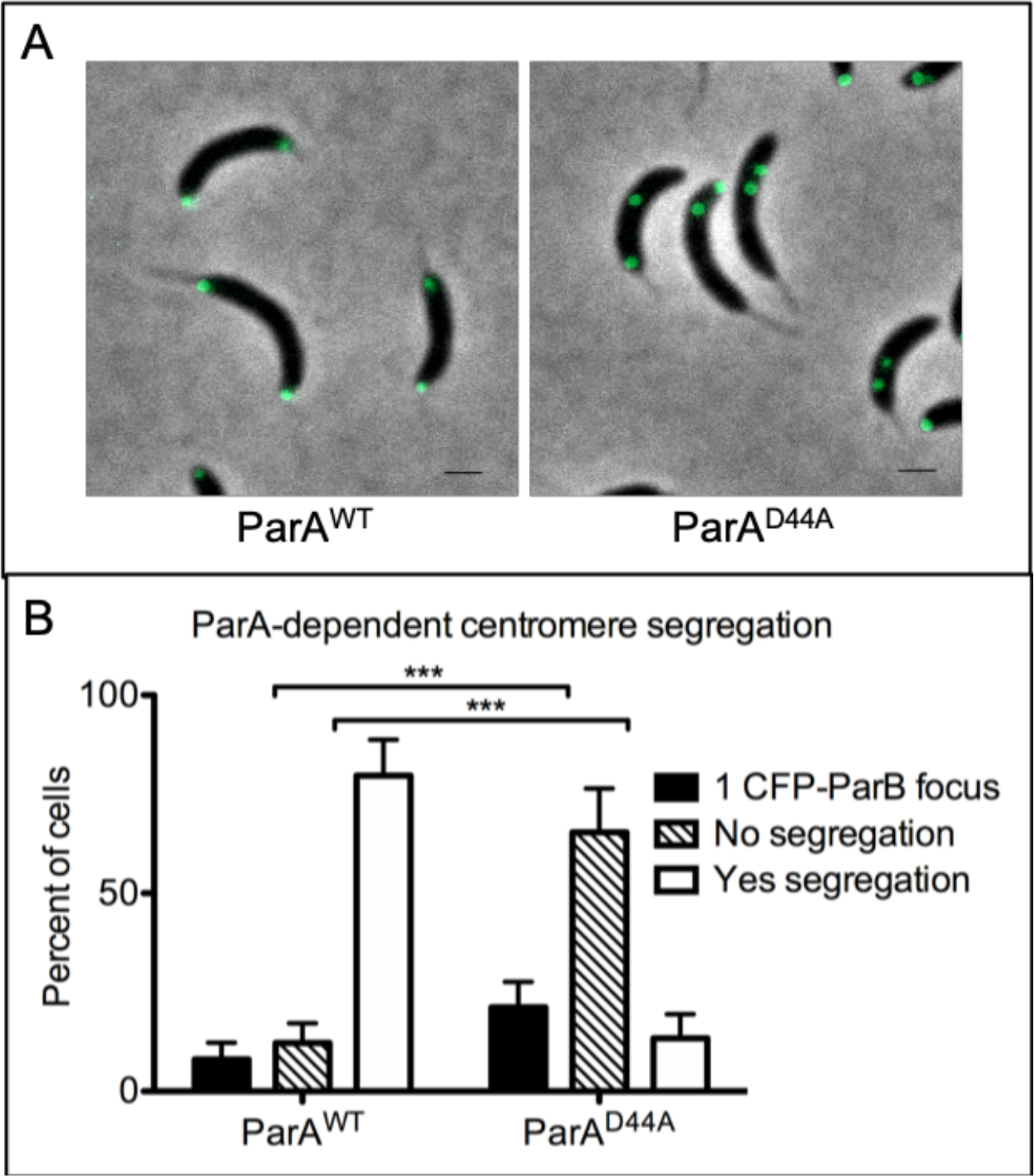
Centromere segregation in the opposite direction requires active ParA. Cells with background *parB*::*cfp-parB*, *dnaA*::Ω, *vanA*::*dnaA* with either ParA^WT^ (PM109) or ParA^D44A^ (PM121) were grown in the absence of vanillate to allow for centromere translocation. After 3 h of DnaA depletion, cultures were supplemented with vanillate (250 μM) to express DnaA and xylose (0.3 %) to induce the expression of ParA^D44A^ variant protein. Cells were imaged by spotting 2 μL of cells on 1 % agarose pads. (A) Phase contrast fluorescence micrographs of PM109 expressing wild-type ParA^WT^ (left) and PM121 expressing ATP hydrolysis variant ParA^D44A^ (right). Scale bar = 1 μm. (B) Frequencies of centromere segregation were quantified based on localization of CFP-ParB (green foci). The data represents analyses of three independent experiments. Bar graph illustrates the mean ± SD values. Statistical analysis: two-way ANOVA, ***P<0.001.

### Re-localization of the centromere locus triggers rearrangement of ParA gradient

In wild-type *C. crescentus*, ParA forms a gradient with high concentrations at the new pole that gradually decrease toward the stalked pole (Fig. 1) [26–28]. Our observation that the centromere could be segregated in the opposite direction suggested that cells with centromeres at the new pole rearranged the gradient of ParA. To determine whether ParA could change the orientation of its gradient, we assessed the localization patterns of ParA using the background of a *parA* merodiploid strain that contains the wild-type allele of *parA* at the native locus and a fluorescently-tagged *parA* (ParA-mCherry) under the inducible promoter for xylose [36]. This construct grown under normal DnaA expression conditions, in the presence of vanillate, displayed wild type localization dynamics of ParA-mCherry. However, cells with un-replicated centromeres that had translocated to the new pole displayed a completely flipped pattern of the ParA-mCherry gradient (high levels at the stalked pole that gradually decrease toward the new pole) (Fig. 6A). Our data revealed that similarly to DnaA and replication initiation, ParA can rearrange to initiate the segregation of the centromere from outside the stalked pole.

**Figure 6.**
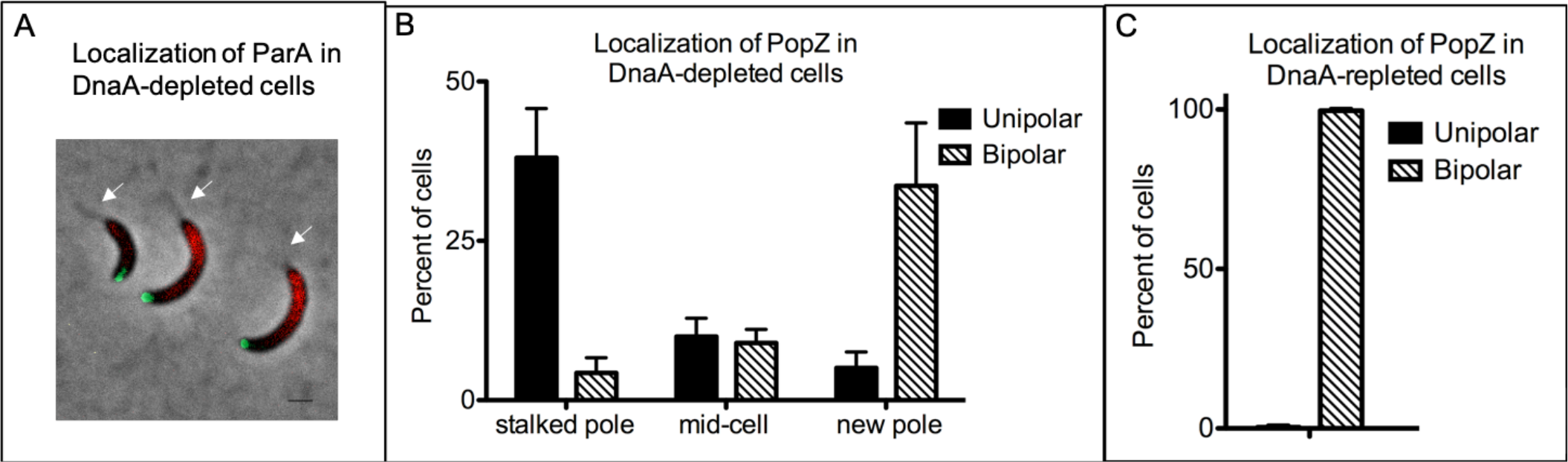
Localization of ParA and PopZ in cells with translocated centromeres. (A) Translocation of centromere to the new pole in DnaA depleted cells results in flipped ParA-mCherry gradient (red). Green foci represent CFP-ParB (centromeres). White arrows indicate location of the stalks. PM503 cells (*parB*::*cfp-parB*, *dnaA*::Ω, *vanA*::*dnaA xylX::parA-mCherry*) were synchronized and depleted of DnaA for 3 h. The culture was supplemented with xylose (0.1 %) for 2 hours during the time of depletion. After DnaA depletion (3 h), 2 μL of cells were mounted on 1 % agarose pad and imaged using phase contrast fluorescence microscopy. Scale bar = 1 μm. (B & C) Quantification of localization of PopZ in cells depleted (B) and repleted (C) of DnaA. PM247 (*parB*::*cfp-parB*, *dnaA*::Ω, *vanA*::*dnaA xylX*::*mCherry-popZ*) were grown in the absence of vanillate (DnaA depletion) for 3 h and then supplemented with vanillate (DnaA repletion) for 1 h. Cultures were supplemented with xylose (0.1%) to induce the expression of mCherry-PopZ for 1 h prior to isolation of swarmer cells. Phase contrast fluorescent micrographs were obtained just before and after 1 hour of the addition of vanillate. The data represents analyses of three independent experiments. Bar graph illustrates the mean ± SD values.

**Figure 7.**
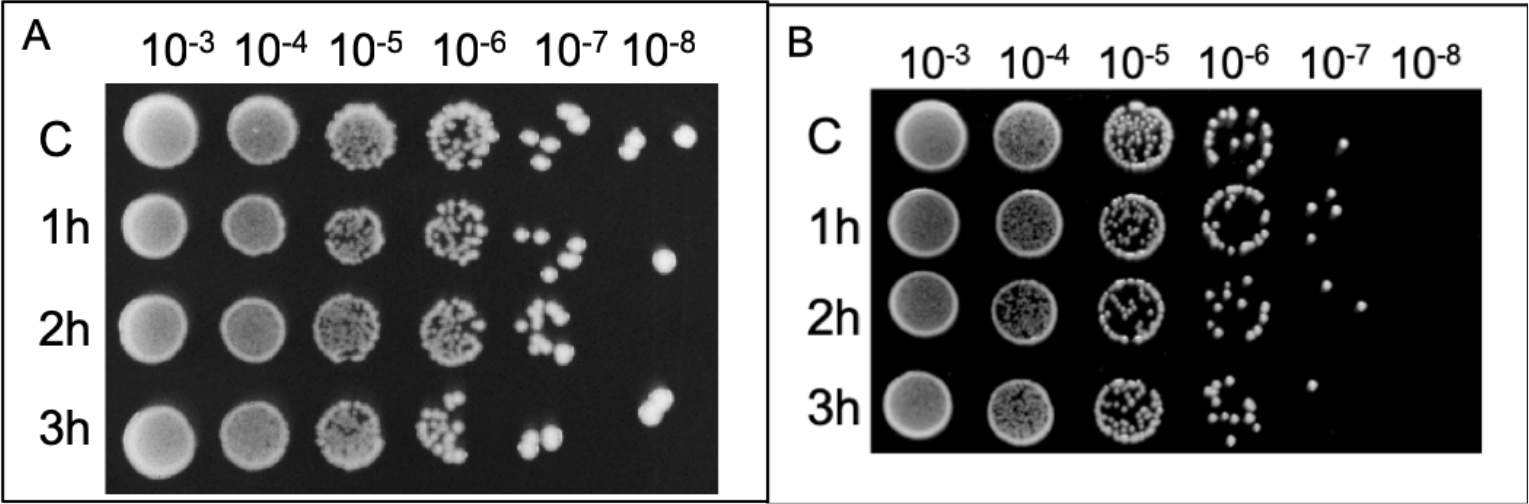
Depletion of DnaA for 3 h does not alter the viability of *C. crescentus*. Figure illustrates colony forming unit (CFU) assays of (A) *parS* (pMT1) *vanA*::*dnaA*, Δ*dnaA*, *xylX*::*cfp-parB* (pMT1; PM500) and (B) *parB*::*cfp-parB*, *dnaA*::Ω, *vanA*::*dnaA* (PM109) cells. The cultures (3 mL) grown up to OD_600_ ~ 0.3 were washed three times with 1X M2 salts as described under methods section and the OD_600_ were set to ~ 0.2 in M2G media (2 mL). DnaA was depleted for 1 h, 2 h and 3 h in separate cultures at 28°C and CFU assay were performed. Control sample (C) were not depleted of DnaA and incubated with vanillate (250 μM) for 3 h. PYE plates were incubated at 28°C for 2 days prior to obtaining the images. The data shown are a representative of three independent replicates.

### PopZ bi-polar localization is triggered independently of chromosome replication

The directionality of ParA’s chromosome segregation from the stalked pole to the new pole has been proposed to be dependent on the localization of the anchoring protein PopZ [36]. To determine whether the onset of chromosome replication/segregation from outside the stalked pole altered PopZ localization dynamics, we tracked the localization of PopZ by using cells expressing a functional fusion protein mCherry-PopZ. In our control experiment with cells grown in the presence of vanillate (the *dnaA* inducer), mCherry-PopZ foci localized one at each pole upon the onset of chromosome replication and segregation [21–23, 31]. In cells with translocated un-replicated centromeres, mCherry-PopZ localization was dependent on the localization of CFP-ParB (Fig. 6B). ~ 90 % of cells with CFP-ParB at the stalked pole displayed a single mCherry-PopZ focus also localized at the stalked pole. ~90 % of cells with CFP-ParB localized at the new pole displayed two PopZ-CFP foci with one localized at each pole. Notably, cells with CFP-ParB localized at mid-cell displayed an equal combination of cells with either one mCherry-PopZ focus localized at the stalked pole or two PopZ-CFP foci localized one at each pole. Upon the induction of chromosome replication by the addition of the inducer vanillate, cells with bi-polar localization of PopZ seemed remained bi-polar (Fig. 6B). Regardless where the centromere was localized when replication initiation was induced by vanillate supplementation, ~100 % of cells displayed bi-polar localization of mCherry-PopZ (Fig. 6C).

### Effects in viability from initiating chromosome replication/segregation from outside the stalked pole

To determine whether initiating chromosome replication and/or segregation from outside the stalked pole altered the viability of *C. crescentus,* we analyzed CFU of cells that had undergone ori/centromere translocation away from the stalked pole in the absence of replication. Cells PM500 (fluorescent label near *ori*) and PM109 (fluorescent label ParB-centromere) were spotted immediately after DnaA depletion time (1 h to 3 h) on plates containing the inducer for *dnaA* expression (supplemented with vanillate). Our CFU analyses revealed no significant differences between cells that initiated chromosome replication/segregation from the new pole (or mid-cell) compared to wild type conditions. Our data suggest that cells can recover relatively quickly after initiating chromosome replication/segregation from outside the stalked pole.

## DISCUSSION

By inducing the movement of *ori* and centromere away from their intrinsic sub-cellular location in a replication-independent manner, we have shown that the molecular machineries involved in chromosome replication and segregation are remarkably flexible. Our data revealed that the activity of DnaA and ParA are not restricted to a single subcellular location inside the cell and can successfully induce chromosome replication and segregation from outside the stalked pole in *C. crescentus*. These results suggest that the set of proteins that regulate the asymmetric organization of chromosome replication and segregation must also rearrange, posing a model where these vital chromosomal loci (*ori* and centromere) play a key role in the localization of regulators that maintain cellular asymmetry. Cells that initiate chromosome replication and segregation from outside the stalked pole display no detectable physiological defects compared to wild type conditions. Thus, our data expose the remarkable ability of the cell to reorganize its chromosome along with the set of protein regulators involved in chromosomal maintenance in a relatively short time.

### DnaA’s activity as replication initiator and cell asymmetry

In the absence of membrane-bound organelles, bacteria rely on proteins organized in gradients to establish cellular polarity and carry out asymmetric functions. One example of such organization is the phosphorylated regulator CtrA (CtrA~P), which binds *ori* and inhibits DnaA from initiating replication at one cell pole [37–39]. In pre-divisional cells, a phospho-signaling relay at the cell poles generates an asymmetric concentration gradient of CtrA~P with the highest levels at the new pole that gradually decrease toward the stalked pole [40]. Alterations to the asymmetric concentration gradient of CtrA~P eliminates the asymmetric regulation of DnaA, resulting in cells initiating replication from the new pole [40]. Our data revealed that DnaA can also trigger replication initiation from the new pole in cells with altered subcellular location of *ori* (Fig. 3). Our results can be explained by the undetectable levels in Western blot assays of CtrA in cells depleted of DnaA [41, 42]. DnaA is a transcriptional regulator of *gcrA,* and GcrA is a transcriptional regulator of *ctrA* [41–43]. Consequently, the expression of *ctrA* is indirectly dependent on the levels of DnaA. Thus, cells with *ori* localized at the new pole with depleted levels of DnaA are likely to have none or minimal CtrA~P gradient that is not sufficient to inhibit replication initiation from either pole.

### Regulation of periodicity of DnaA’s activity

Most of the regulators of DnaA’s activity that have been identified so far are negative regulators that prevent the over-initiation of chromosome replication. However, positive regulators that trigger DnaA to initiate replication with such efficient periodicity remain limited. This periodicity of DnaA’s activity is maintained even in *E. coli* cells that were artificially designed to have two spatially separated *oris.* In those cells, DnaA productively initiated replication synchronously from both *ori*s [44]. In *E. coli* and *H. pylori*, a recruiter of DnaA has been characterized to promote the assembly of DnaA’s polymer at *ori* [8–10]. Notably, constitutive expression of *dnaA* has been shown to have no effect on the periodicity of DnaA’s activity suggesting that *dnaA* transcriptional regulation is not the principal modulator of DnaA’s periodicity [18]. In *C. crescentus*, CtrA regulates the spatial activity of DnaA so that chromosome replication only initiates in the stalked cells [18, 38, 39]. However, CtrA is not involved in the periodic activity of DnaA [18]. Thus, the molecular mechanism that triggers DnaA to initiate chromosome replication with such precise periodicity remains unclear. A hypothetical scenario is that some type of regulator that facilitates this periodicity process is found within the microenvironment of where *ori* is localized at the time of replication initiation. Our data advocates that there is no regulator/modulator of DnaA that is fixed at the stalked pole of *C. crescentus*. We cannot however exclude the possibility that a potential modulator does exist, and this modulator could migrate along with *ori* because it either binds directly to *ori* or to DnaA. More work characterizing details about DnaA’s activity at *ori* is required to identify the potential modulator of periodicity or the mechanism that DnaA uses to regulate its temporal activity with such remarkable accuracy.

### Localization of the centromere dictates the orientation of ParA’s activity

The partitioning protein ParA is another example of a protein in bacteria organized in gradients to establish cellular polarity. In *C. crescentus*, ParA forms a gradient with concentrations gradually decreasing from the new pole to the stalked pole [26–28]. Notably, this stable gradient of ParA is established well before chromosome replication and segregation are initiated [26–28]. Thus, the question remains as to what activates this asymmetric organization of ParA. Our data revealed that the organization of the ParA gradient can be completely reconstructed in the opposite orientation by rearranging the location of the centromere (Fig. 6A). We show that ParA can successfully segregate the centromere from the new pole to the stalked pole, which is the reverse direction to its wild type mechanism. We propose that once a stable gradient of ParA is formed in cells with translocated un-replicated-centromeres, the ParA-DNA interaction relay previously shown to provide the force necessary for centromere segregation [45] can initiate and segregate one centromere in the reverse direction.

### Robustness of cells to re-organize

The view that bacteria are simply a bag of enzymes with no level of organization is long gone. Instead, the appreciation of the high levels of organization that bacteria manage in such limited spaces is mind blowing. In *C. crescentus*, an array of signaling proteins localize differentially at each pole. This organization orchestrates the progression of the cell cycle (i.e. chromosome replication, segregation, cytokinesis) with the development of the stalked and new pole. The centromere in *C. crescentus* serves as a hub of proteins involved in multiple important functions involved polar development. In this work, we asked what happens to the ability of cells to grow when the organization of the two key chromosomal loci (*ori* and centromere) are flipped in orientation. We first showed that the regulators (DnaA and ParA) can easily follow the new location of these sites and proceed with their activities, and in the case of ParA proceed segregation in the reverse direction. Notably, cells were able to recover the “forced” rearrangement of these chromosomal loci and continue to grow with no measurable delays. Our results revealed the robustness and flexibility that cells have to rearrange and reorganize the polar organization proteins that interact with *ori* and centromere.

## MATERIALS AND METHODS

### Bacterial strains and growth conditions

Wild-type strain CB15N (NA1000) was used to generate all *C. crescentus* derivatives (Table 1) in this study. Cells were grown in either rich (PYE) or minimal (M2G) medium [46] from freezer stock at 28 °C. Growth medium was supplemented with vanillate (Sigma-Aldrich, 250 μM) and/or 0.3 % D−(+)−xylose (Sigma-Aldrich) as specified. When noted, the liquid/solid media was supplemented with appropriate antibiotics; for *C. crescentus* kanamycin 5/25 μg/mL, streptomycin 5/5 μg/mL, tetracycline 1/2 μg/mL, and for *E. coli* kanamycin 30/50 mg/mL.

**Table 1.** List of strains and plasmids used in this study.

| Strain or plasmid | Description | Source or reference |
| --- | --- | --- |
| **C. crescentus strains** |  |  |
| PM1 | Wild-type CB15N (NA1000) | Evinger M and Agabian N, 1977 |
| PM109 | CB15N, $\Delta vanA$, *parB::cfp-parB*, *dnaA::*$\Omega$ (*strp<sup>R</sup>/spec<sup>R</sup>*), *vanA::dnaA* | Mera *et al*, 2014 |
| PM121 | *xylX::parA(D44A)-yfp* (*tet<sup>R</sup>*) in PM109 (*strp<sup>R</sup>*) | This study |
| PM247 | *xylX::mCherry-PopZ* (*tet<sup>R</sup>*) in PM109 (*strp<sup>R</sup>*) | Mera *et al*, 2014 |
| PM433 | CB15N, pMT1 *parS* | This study |
| PM438 | CB15N, pMT1 *parS*, *xylX::yfp-parB* (pMT1) (*kan<sup>R</sup>*) | This study |
| PM500 | CB15N *parS* (pMT1), *vanA::dnaA*, $\Delta dnaA$, *xylX::cfp-parB* (pMT1) (*kan<sup>R</sup>*) | This study |
| PM503 | pDNA219 (*parA-mCherry* under xylose promoter) (*kan<sup>R</sup>*) in PM109 (*strp<sup>R</sup>*) | This study |
| **Plasmids** |  |  |
| pNPTS138 | Nonreplicating vector for allelic replacement (*kan<sup>R</sup>*) *oriT sacB* | Alley M. R. K., unpublished data |
| pXCHYC-2 | Integrating plasmid for xylose inducible expression of mCherry tagged CCNA_03869 (*parA*); (*kan<sup>R</sup>*) | Thanbichler M. *et al*, 2007 |
| pXCFPN-2 | Integrating plasmid for xylose inducible expression of CFP tagged *parB*(pMT1) (*kan<sup>R</sup>*) | Thanbichler M. *et al*, 2007 |
| pDNA214 | pNPTS138- *parS* (pMT1) sequence flanked by 600bp UP/DWN at base 1108 for insertion of *parS* (pMT1) near *ori*. | This study |
| pDNA216 | pXCFPN-2 – *xylX::cfp-parB* (pMT1) (*kan<sup>R</sup>*) | This study |
| pDNA217 | pNPTS138- *dnaA* sequence flanked by 600bp UP/DWN of CCNA_02476. To replace *vanA* with *dnaA* | This study |
| pDNA218 | pNPTS138- 600bp UP/DWN CCNA_00008 for *dnaA* deletion | This study |
| pJP49 | pXYFPC5-*parA*(D44A) | Ptacin J.L. *et al*, 2010 |
| pDNA219 | CCNA_03869 cloned into pXCHYC-2 at NdeI 5' and SacI 3' restriction enzyme sites | This study |
| pMS138 | pMCS4 – *parS* (pMT1) | Schwartz M. & Shapiro L., 2011 |
| pMS139 | pXYFPN-2 - *xylX::yfp-parB*( pMT1) (*kan<sup>R</sup>*) | Schwartz M. & Shapiro L., 2011 |

Plasmids constructed in this study were created by cloning PCR products amplified using wild-type CB15N (NA1000) genomic DNA or *Yersinia pestis* KIM5 pMT1 DNA into pNPTS138, pXCHYC-2 and pXCFPN-2 [33] vectors. The constructs were transformed into *E. coli* DH5α cells and grown at 37 °C in Luria-Bertani (LB) medium. All primers used for cloning are listed in Table 2. Plasmid carrying CFP-pMT1 *parB* was done by the pMT1 *parB* gene sequence isolation with KpnI and NheI restriction from PM396 (LS5269) and ligation to the equally treated xylose-inducible integrating plasmid pXCFPN-2 (kan^R^) [24]. *parA* gene was cloned into integrating pXCHYC-2 (kan^R^) plasmid under a xylose inducible promoter to express mCherry tagged C-terminal protein fusions [33]. Gibson cloning method [47] was used to construct the plasmids used to delete or insert a gene into the *C. crescentus* genome. To insert pMT1 *parS* site approximately 1kb far away from the *ori*, the cloned pMT1 *parS* sequence from PM395 (LS5270) and around 600bp of CCNA0001 C-terminus and CCNA0002 N-terminus sequences were assembled into pNPTS138 plasmid [24]. The plasmid used for *vanA*::*dnaA* substitution was accomplished by assembling into pNPTS138 plasmid the cloned *dnaA* gene sequence and 600 bp of the flanking upstream and downstream sequences of *vanA*. For dnaA deletion purposes, 600 bp of flanking upstream and downstream sequences of *dnaA* were cloned into pNPTS138 plasmid.

**Table 2.**
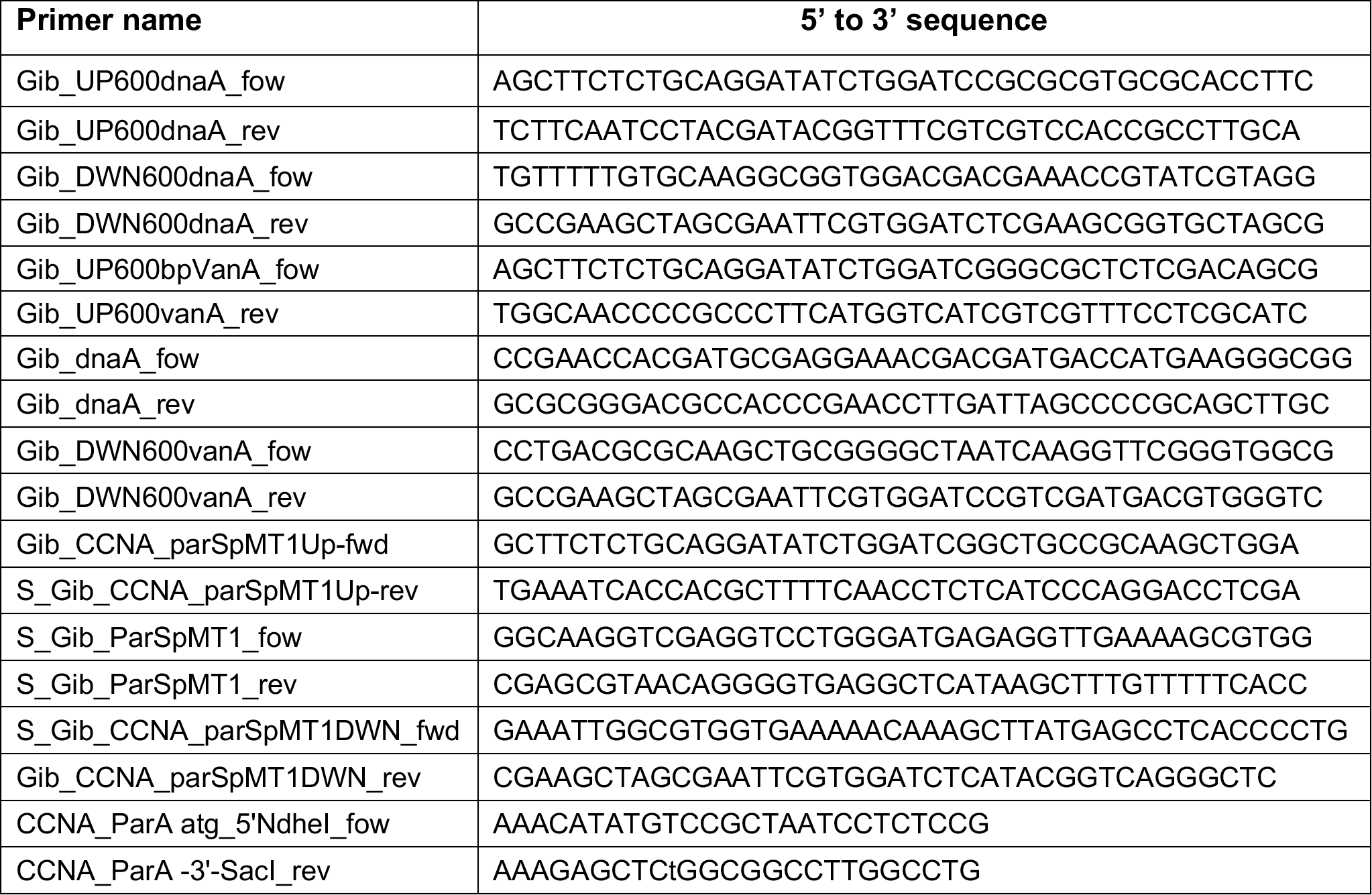
List of primers used in this study. Primers that start with “Gib” were used in Gibson cloning technique [47].

### Growth assays

Overnight cultures grown from *C. crescentus* frozen stocks in M2G liquid media were diluted to 0.2 OD_600nm_ (2mL) in 13 mm glass tubes. Culture tubes were incubated at 28 °C and the optical density at OD_600nm_ was monitored every hour to monitor the growth of bacteria. Rate of bacterial growth was calculated using the exponent of the growth curves on semi-log plots.

### Synchrony

A culture of *C. crescentus* in M2G (15 mL) was inoculated with a saturated overnight M2G culture and was grown up to OD_600_ around 0.3. Media was supplemented with vanillate (250 μM) and antibiotics as noted. Cells were pelleted using centrifugation at 6000 rpm for 10 min at 4 °C. The cell pellet was resuspended in about 800 μL of 1x M2 salts and mixed well with percoll (900 μL, Sigma Aldrich). Swarmer cells (bottom layer) were collected and separated out from the stalked/predivisional cells (top layer) using percoll density gradient by centrifuging at 11,000 rpm for 20 min at 4 °C. The cells were washed twice with cold 1× M2 salts by spinning at 8000 rpm for 3 min at 4 °C and resuspended in M2G media to the appropriate OD_600_. When cells were not synchronized, the cultures grown to around 0.3 OD_600_ were pelleted and washed with 1× M2 salts three times.

### Fluorescence microscopy

The cells (1-3 μL) were spotted on agar pads (1 % agarose in M2G) and imaged using phase contrast and fluorescent microscopy in Zeiss Axio Observer 2.1 inverted microscope, set up with a Plan-Apochromat 100×/1.40 Oil Ph3 M27 (WD=0.17 mm) objective, AxioCam 506 mono camera and ZEN lite software. Agar pads supplemented with vanillate (250 μM) were used in time-lapse assays when needed. Images were analyzed using ImageJ software (https://imagej.nih.gov/ij/docs/menus/analyze.html) [48] and localization of fluorescent foci was counted using the Cell Counter plugin.

### Colony forming units (CFU) assay

Serial dilutions of culture in ten-fold were done by mixing 10 μL of culture with 90 μL of PYE medium in a sterile 96 well plate. 5 μL of each sample was spotted on PYE agar (1.5 %) plates supplemented with vanillate (250 μM) if needed. CFU counts were obtained from the plates incubated at 28 °C for two days.

## AUTHOR CONTRIBUTIONS

ABM, IPM, and PEM designed the study, acquired/analyzed data, and wrote the manuscript.

## Supporting information

Supplemental Material Melendez

