## Supplemental Material Melendez for "Chromosome dynamics in bacteria: triggering replication at opposite location and segregation in opposite direction"

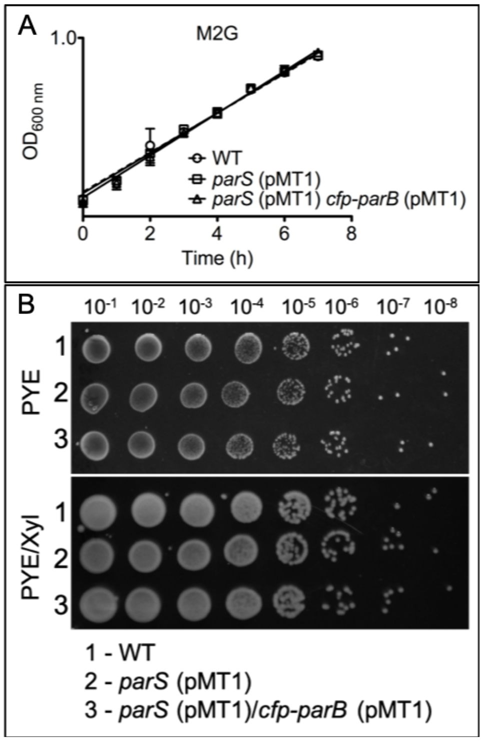


**Fig. S1.** Construction of *Y. pestis parS*/*cfp-parB* (pMT1) system into *C. crescentus* near *ori* does not alter cell viability. (A) Exponential state growth curves (n = 3) of *C. crescentus* WT (PM1: NA1000, wild-type), *parS* (pMT1) (PM433: cells with *Y. pestis parS* sequence inserted near *ori*), and *parS*/*cfp-parB* (pMT1) (PM438: PM433 with CFP-ParB from *Y. pestis* expressed from xylose inducible promoter). Cells were grown in minimal media (M2G). The cultures (2 mL) were set to $OD_{600}$ ~ 0.2 and the growth was monitored by measuring the absorbance at 600 nm every hour. Expression of *cfp-parB* (pMT1) was induced by adding xylose (0.3%). (B) The colony forming units (CFU) assay of *C. crescentus* wild-type (WT, PM1), (PM433) *parS* (pMT1) and (PM438) *parS*/*cfp-parB* (pMT1) were spotted on PYE plates in the presence or absence of xylose (0.3%). CFU assays were carried out as described in Material and Methods. Cultures were grown at 28 °C and the CFU plates were incubated for 2 days before imaging. Data shown are representative of three independent replicates.

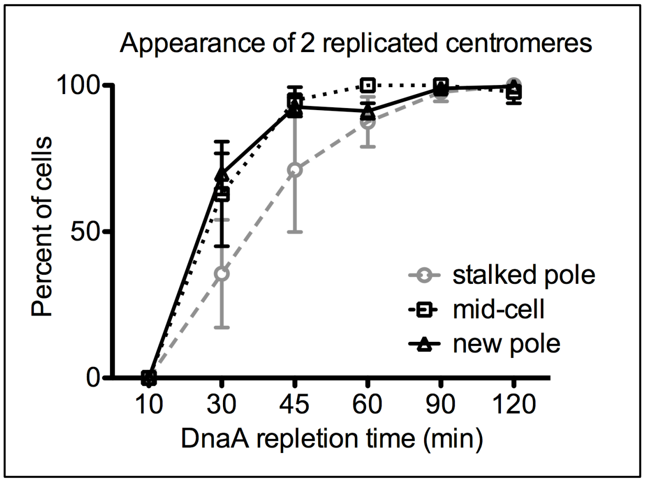


**Fig. S2.** Frequencies of chromosome replication initiation based on centromere localization. Plotted are the percent of *C. crescentus* cells (*parB*::*cfp-parB*, *dnaA*::Ω, *vanA*::*dnaA*; PM109) with 2 CFP-ParB foci based on the initial localization of *parS* (8 kb away from *ori*). DnaA was depleted for three hours in swarmer cells isolated from mini-synchrony. Vanillate (250 μM) was added to the media (time 0 min) and the cells (2 μL) were spotted immediately on an agarose pad (1% in M2G media) supplemented with vanillate (250 μM). Time lapse images were obtained using phase contrast microscopy at the given time intervals and the appearance of 2 replicated centromeres were monitored. Data represent the mean with standard deviation (SD) of three independent replicates. Comparison of the two-way ANOVA in-between the frequencies of replication initiation at or near stalked pole and mid-cell or new pole were significantly different at 30-min and 45-min time points (stalked pole and mid-cell *$p<0.05$, stalked pole and new pole ***$p<0.001$ at 30-min and *$p<0.05$ at 45-min).

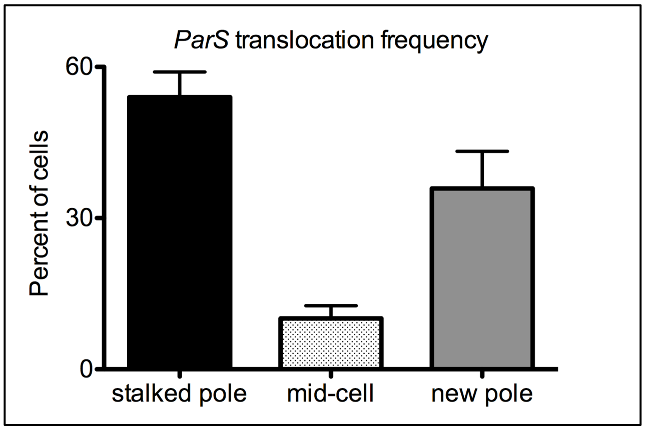


**Fig. S3.** Frequencies of DnaA-dependent replication-independent centromere translocation. Localization of CFP-ParB foci (centromere) at or near stalked pole, mid-cell, and at or near new pole of *C. crescentus* cells were calculated in DnaA depleted cells (*parB*::*cfp-parB*, *dnaA*::Ω, *vanA*::*dnaA*; PM109) for 3 hours. Swarmer cells were collected by synchrony and DnaA was depleted following the protocol described elsewhere [Mera *et al.* 2014]. Data represent the mean ± SD (standard deviation) of three independent replicates.
